# Kinetics of HTLV-1 reactivation from latency quantified by single-molecule RNA FISH and stochastic modelling

**DOI:** 10.1101/631697

**Authors:** Michi Miura, Supravat Dey, Saumya Ramanayake, Abhyudai Singh, David Rueda, Charles R. M. Bangham

## Abstract

The human T cell leukemia virus HTLV-1 establishes a persistent infection *in vivo* in which the viral sense-strand transcription is usually silent at a given time in each cell. However, cellular stress responses trigger the reactivation of HTLV-1, enabling the virus to transmit to a new host cell. Using single-molecule RNA FISH, we measured the kinetics of the HTLV-1 transcriptional reactivation in peripheral blood mononuclear cells (PBMCs) isolated from HTLV-1^+^ individuals. The abundance of the HTLV-1 sense and antisense transcripts was quantified hourly during incubation of the HTLV-1-infected PBMCs *ex vivo*. We found that, in each cell, the sense-strand transcription occurs in two distinct phases: the initial low-rate transcription is followed by a phase of rapid transcription. The onset of transcription peaked between 1 and 3 hours after the start of *in vitro* incubation. The variance in the transcription intensity was similar in polyclonal HTLV-1^+^ PBMCs (with tens of thousands of distinct provirus insertion sites), and in samples with a single dominant HTLV-1^+^ clone. A stochastic simulation model was developed to estimate the parameters of HTLV-1 proviral transcription kinetics. In PBMCs from a leukemic subject with one dominant T-cell clone, the model indicated that the average duration of HTLV-1 sense-strand activation by Tax (i.e. the rapid transcription) was less than one hour. HTLV-1 antisense transcription was stable during reactivation of the sense-strand. The antisense transcript *HBZ* was produced at an average rate of x~0.1 molecules per hour per HTLV-1^+^ cell; however, between 20% and 70% of HTLV-1-infected cells were *HBZ*-negative at a given time, the percentage depending on the individual subject. HTLV-1-infected cells are exposed to a range of stresses when they are drawn from the host, which initiate the viral reactivation. We conclude that whereas antisense-strand transcription is stable throughout the stress response, the HTLV-1 sense-strand reactivation is highly heterogeneous and occurs in short, self-terminating bursts.

**Author summary:** Human retroviruses such as HIV-1 and HTLV-1 (human T cell leukemia virus) can establish a latent infection in the host cell. However, these viruses need to be able to produce viral genome to propagate in a new host. HTLV-1-infected cells are transmitted through breastfeeding, blood transfusion and sexual contact, and HTLV-1 restores transcription once the infected cells are drawn from infected individuals. We measured the kinetics of the HTLV-1 transcriptional reactivation in blood cells isolated from HTLV-1^+^ individuals by single-molecule RNA FISH. Viral transcripts were visualised as diffraction-limited spots and their abundance was quantified at one-hour intervals. The onset of the virus transcription peaked after one to three hours of incubation. In each cell, a short period of slow HTLV-1 transcription was followed by a phase of rapid transcription. Computer simulation, based on experimental data on PBMCs from a leukemic patient with a single dominant HTLV-1-infected T cell clone, indicated that this rapid transcription from the HTLV-1 sense-strand promoter activated by Tax was terminated in less than an hour. The HTLV-1 antisense transcript *HBZ* was constantly produced at a low level, and 50% ± 20% of HTLV-1^+^ cells were negative for *HBZ* at a given time. These results demonstrate how rapidly HTLV-1 is reactivated and potentially becomes infectious, once HTLV-1^+^ cells are transmitted into a new host.

## Introduction

Retroviral latency constitutes the main barrier to eradicating the infection from the host. The provirus is not completely silent, but is intermittently transcribed to produce viral particles that infect new clones of cells [1]. This intermittent transcription is partly due to the apparently stochastic nature of gene regulation [2]. However, it is poorly understood what regulates the balance between retroviral latency and transcription. Attempts to reactivate HIV-1 to expose it to drug therapy and the immune response - the shock-and-kill strategy [3] - have not been successful so far because it has not been possible to reactivate all the HIV-1 provirus in the reservoir [4]. Therefore, it is essential to understand the mechanisms and the extent of heterogeneity in the viral reactivation. Human T cell leukemia virus type 1 (HTLV-1) is a pathogenic retrovirus, which causes fatal and disabling diseases [1]. HTLV-1 remains largely latent *in vivo*, but undergoes transcriptional reactivation once the HTLV-1-infected lymphocytes are drawn from the circulation [5]. Here, we exploit this pattern of predominant HTLV-1 latency *in vivo* and spontaneous transcriptional reactivation *ex vivo* to study the kinetics of retroviral reactivation.

It is estimated that 10 million people across the globe are living with HTLV-1 [6]. HTLV-1 resides mainly in CD4^+^ T lymphocytes, and causes adult T cell leukemia/lymphoma (ATL), a CD4^+^ T cell malignancy, in some 5% of HTLV-1-infected individuals. HTLV-1 also causes, in up to another 5%, HTLV-1-associated myelopathy/tropical spastic paraparesis (HAM/TSP; hereafter abbreviated to HAM) and other inflammatory disorders [1]. ATL typically presents as a monoclonal expansion of HTLV-1-infected T lymphocytes that is characterized by a unique integration site of the HTLV-1 provirus in the host genome [7], with an underlying polyclonal population of lower abundance. By contrast, patients with HAM and asymptomatic HTLV-1 carriers have a highly polyclonal population of ~10^4^ HTLV-1^+^ T cell clones [7, 8].

HTLV-1 transcribes both the sense and antisense strand of its 9 kb provirus. The long terminal repeats (LTRs) located at the ends of the provirus serve as the promoters for sense and antisense transcription respectively. HTLV-1 Tax, produced from the sense strand, is a transcriptional transactivator protein that triggers NF-κB activation and other transcriptional cascades as well as its own promoter at the LTR (reviewed in ref [9]), exerting a strong positive feedback on the sense-strand transcription of the provirus. The HTLV-1 transcript *tax* is undetectable *in vitro* in freshly isolated cells from most HTLV-1-infected individuals [10]. However, the presence of chronically activated cytotoxic T lymphocytes (CTLs) that recognize sense-strand-encoded products, particularly the highly immunodominant Tax protein [11], indicates that the sense strand is intermittently expressed *in vivo*. By contrast, transcripts of the HTLV-1 bZIP factor (*HBZ*) from the antisense strand are usually detectable *in vitro* at the cell population level in peripheral blood mononuclear cells (PBMCs) obtained from HTLV-1-infected individuals [10].

In a previous study, we applied single-molecule RNA FISH (smFISH) [12, 13] to quantify HTLV-1 sense and antisense transcription in HTLV-1^+^ T cell clones isolated from HTLV-1-infected individuals by limiting dilution and maintained *in vitro* [14]. We observed that the HTLV-1 sense strand is transcribed in an occasional yet intense transcriptional burst. The antisense transcript *HBZ*, however, was much more constantly expressed, albeit at a low level.

To propagate between cells and between individuals, HTLV-1 must restore the sense-strand transcription from latency and produce viral components to form a virological synapse at the site of cell-to-cell contact [15]. The infection spreads to a new host when HTLV-1-infected T lymphocytes are transmitted by breastfeeding, sexual contact and blood transfusion. HTLV-1 sense transcripts rapidly become detectable when the HTLV-1-infected T lymphocytes are drawn from the circulation: the transcriptional reactivation is mediated by p38 MAP kinase activation [5] in response to diverse forms of cell stress. In the present study, we quantified this transient HTLV-1 transcriptional reactivation using smFISH [14] and used stochastic simulation to derive numerical estimates of the parameters of HTLV-1 reactivation from latency.

## Results

### HTLV-1 sense-strand shows two distinct phases of transcription in *ex vivo* culture

HTLV-1-infected PBMCs carry a single copy of the provirus in the genome. HTLV-1 rapidly initiates transcriptional reactivation of the sense strand once the PBMCs are introduced into culture. We used smFISH to quantify the frequency and intensity of HTLV-1 reactivation at the single-cell level (Fig 1A). We removed CD8^+^ cells to avoid CTL-mediated killing of HTLV-1-infected PBMCs that express sense-strand antigens (particularly Tax) during culture [16]. The PBMCs were sampled every hour and subjected to smFISH.

**Fig 1.**
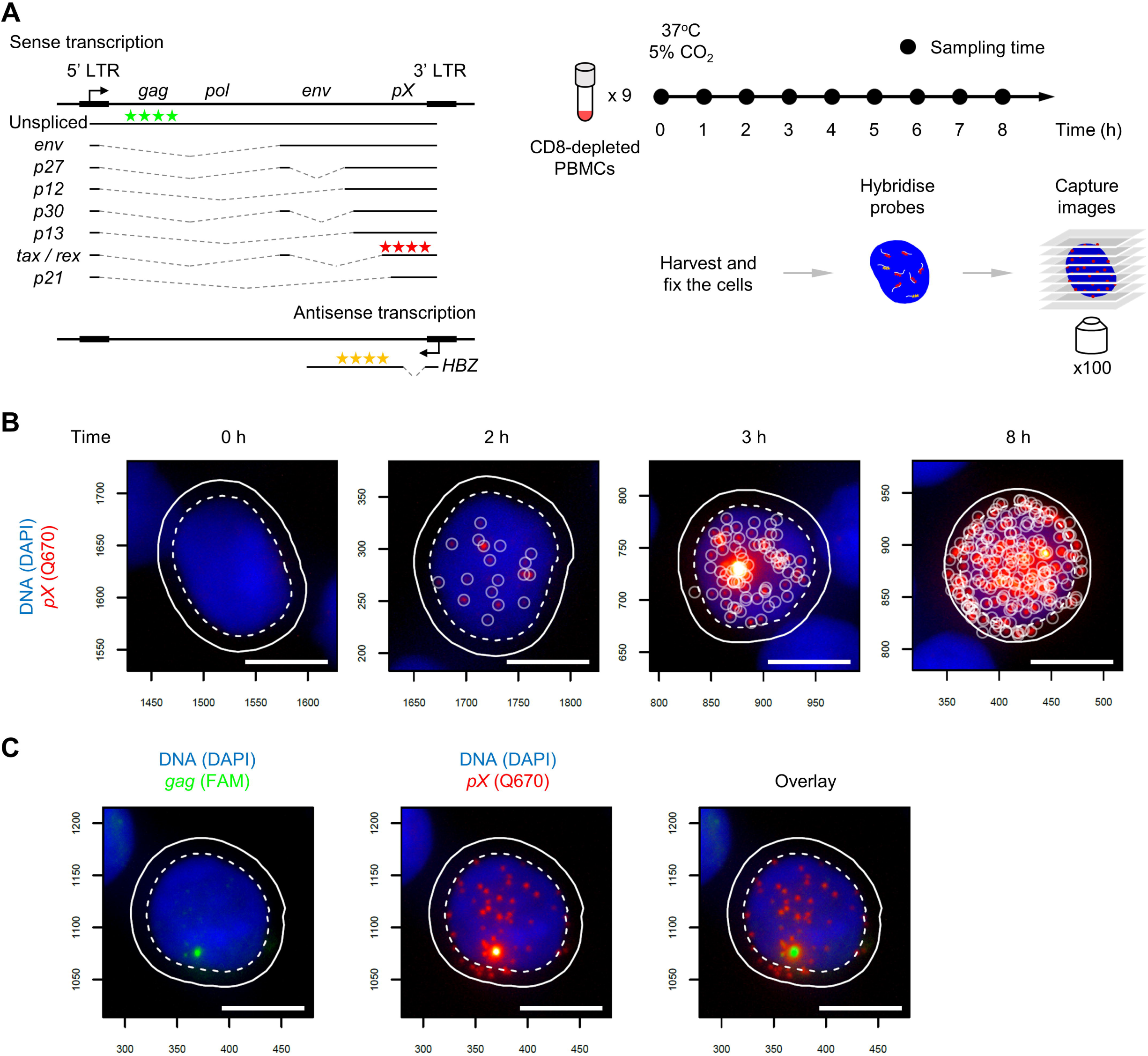
Intense transcription of the HTLV-1 sense strand in PBMCs identified by smFISH. (A) HTLV-1 transcripts, smFISH probes and sample preparation for smFISH. Three sets of probes for smFISH are indicated that hybridize to HTLV-1 transcripts: sense-strand in the *pX* region (Q670, red stars), *HBZ* (Q570, yellow stars), and unspliced sense transcripts containing *gag* region (FAM, green stars). (B) Representative images of HTLV-1^+^ PBMCs with the sense-strand transcripts at the indicated time points. Blue area indicates the nucleus stained with DAPI, and red spots indicate the HTLV-1 sense transcripts. Scale bar (white) = 5 μm. (C) A site of intense transcription identified by the probes for *gag*. The image on the left shows the *gag* staining (green). The image overlaid with sense transcript spots (red, shown in the middle) is presented on the right. The cell presented is from a HAM patient coded TDZ sampled at 4 hours of incubation.

The sequence of representative images of cells (from a patient with HAM, coded TDZ) in Fig 1B shows the increase in the abundance of HTLV-1 sense transcripts over time. Sense-strand transcripts were virtually absent at 0 h (i.e. without incubation), and became detectable in a fraction of cells after 2 h of incubation. Subsequently, a bright spot appeared in the nucleus in those cells in which the sense-strand transcripts were present. The bright spot is an indication of rapid transcription [17], where the transcripts are produced faster than they diffuse away from the site of transcription, resulting in the accumulation of mRNAs at the position of the HTLV-1 provirus in the nucleus. This site contains nascent, unspliced transcripts, as revealed by an additional set of probes that targets *gag* (Fig 1C) which is otherwise spliced out of the sense transcripts (Fig 1A), indicating that the rate of transcription is even faster than that of the splicing reaction [12]. In this paper, we define a transcriptional burst in a single cell as the presence of viral transcripts in the nucleus [18, 19], as opposed to the rapid increase in the abundance of HTLV-1 transcripts over time at the population level. The occurrence of the intense spot in the nucleus was typically associated with the presence of ~10 to 20 sense transcripts in the nucleus. This observation is consistent with the idea that the positive feedback exerted by Tax protein on the sense-strand promoter occurs after a short interval following the first appearance of sense-strand transcripts, during which *tax* mRNA enters the cytoplasm, is translated, and Tax protein re-enters the nucleus. Thereafter, we observed cells carrying a large number of sense-strand transcripts - up to 300 molecules (Fig 1B).

### Cell-to-cell variation in the intensity of HTLV-1 sense-strand reactivation in HTLV-1^+^ PBMCs

The population of PBMCs imaged by smFISH contained both HTLV-1-infected and uninfected cells. We wished to analyse the distribution of the HTLV-1 mRNA count within the HTLV-1-infected population. We used the proviral load to calculate the number of HTLV-1-infected cells in the total population (Fig 2A).

**Fig 2.**
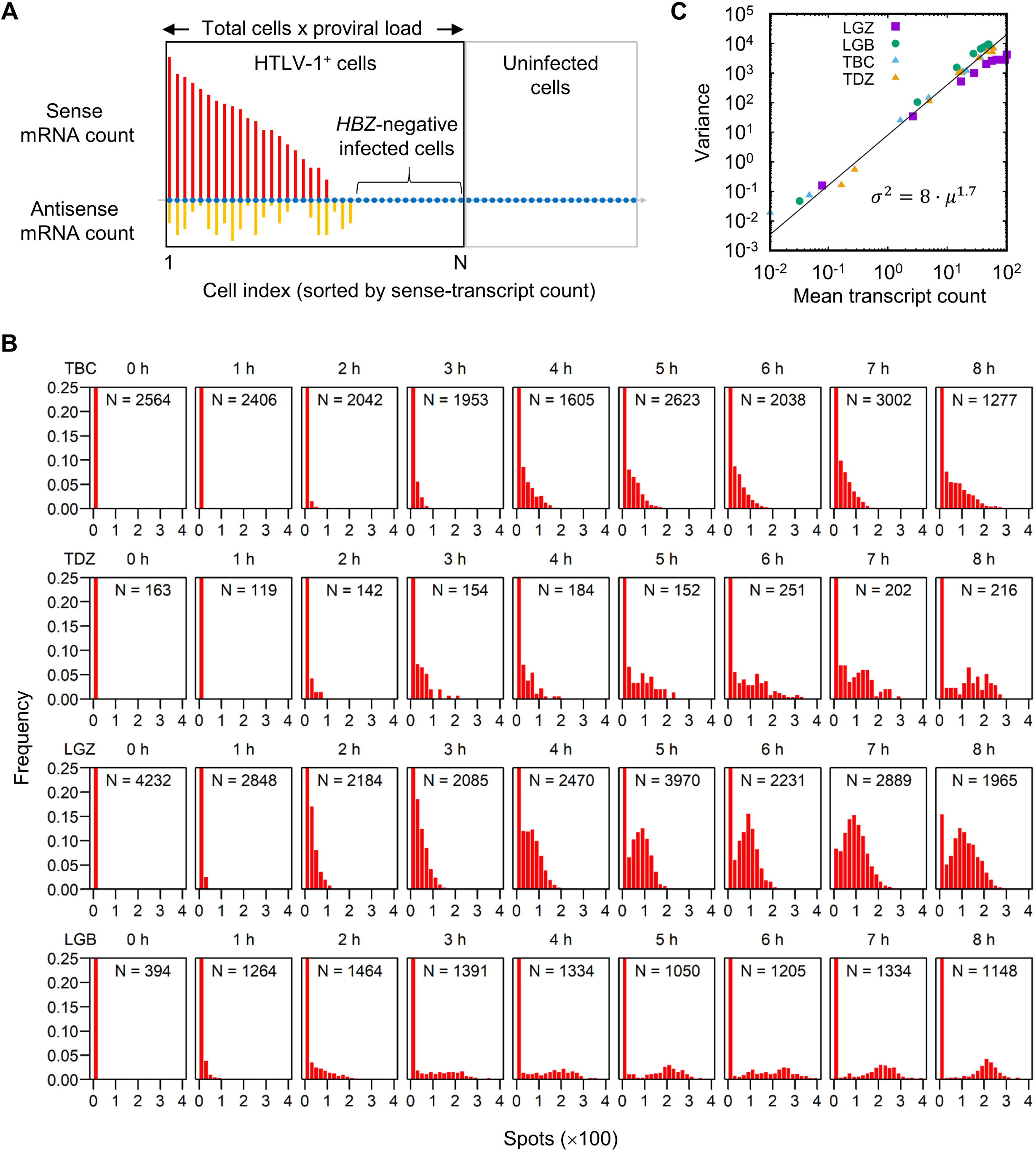
Sense transcript count per HTLV-1^+^ cell: progression over time. (A) Schematic diagram of the identification of the HTLV-1^+^ population for analysis. The entire population (in the black and grey rectangles) are the cells identified and analysed by microscopy. The size of the HTLV-1-infected population (black rectangle) was estimated by multiplying the total cell count by the proviral load as measured by ddPCR. Also see Materials and Methods. (B) Sense-transcript count per HTLV-1^+^ cell (bin size = 20, note the units) at each time point for the two patients with HAM (TBC and TDZ) and two with ATL (LGZ and LGB). Note that the first few bins in each case are out of scale on the y-axis. In the inset for each histogram is indicated the number of cells analysed. (C) The mean and variance of the sense transcript count per HTLV-1^+^ cell obtained from four patients, each containing nine time points.

The distribution of the number of sense-strand transcripts per HTLV-1^+^ cell over time is shown in Fig 2B. In each HTLV-1^+^ subject examined (TBC, TDZ, LGZ and LGB), the total number of transcripts increased continuously during the 8 hours of incubation. The distribution in the sample from one patient with HAM (TBC) presented a long tail of cells that carried a larger number of sense transcripts (up to 250 molecules), with a continuous distribution of cells that carried fewer than 50 transcripts. By contrast, a single mode was identified at around 100 molecules in an ATL patient (LGZ) carrying a single dominant clone.

The average count of the HTLV-1 sense-strand transcripts in the top 25% of all the infected cells was 100 to 300 molecules (S1 Fig). A steeper increase in the count was observed between 1 and 2 hours than in the first hour, which accords with the notion that the HTLV-1 sense-strand expression is biphasic.

We postulated that the polyclonal structure of HTLV-1-infected T cell clones in subjects with non-malignant HTLV-1 infection [7] explains the observed diversity in the distribution of the number of sense-strand transcripts per cell, whereas ATL patients with a monoclonal expansion of an HTLV-1^+^ T cell clone would show more uniform distribution of the number of sense transcripts. We plotted the variance vs. the mean of the observed transcript count in the four HTLV-1^+^ subjects with observations at nine time points (Fig 2C). The mean and variance from all the four samples showed a uniform power-law relationship with the exponent 1.7, suggesting that there is no fundamental qualitative difference in the kinetics of transcription between the samples examined.

### HTLV-1 reactivation peaks after a few hours of *ex vivo* culture

To further characterize the transcriptional reactivation of the HTLV-1 sense strand, we quantified the fraction of cells that had become positive for sense transcripts at a given time during short-term culture of PBMCs (Fig 3A). Cells were defined as positive for sense transcripts when they carried 3 or more spots, to discount spurious spots due to noise (S2 Fig; see Materials and Methods). At 0 h, none of the HTLV-1^+^ cells from each of the four HTLV-1^+^ subjects examined (TBC, TDZ, LGZ and LGB) were positive for sense-strand transcripts. After 8 h of incubation, the fraction of positive cells rose to between 0.2 and 0.8, depending on the HTLV-1^+^ subject examined. We assumed that the waiting time until we observed 3 or more molecules in a cell is an accumulation of single stochastic processes, and therefore fitted the plots presented in Fig 3A with a cumulative gamma distribution [20]. The probability distribution function derived from the fitted curve (Fig 3B) shows that the onset of sense-strand transcriptional reactivation peaked within 1 to 3 hours after the start of *ex vivo* culture. The timing of onset was more broadly distributed in samples from patients with HAM (TBC and TDZ) than from those with ATL (LGZ and LGB): the 75% quantiles of the probability distribution were 3.9 h and 4.9 h for TBC and TDZ respectively, whereas those for LGZ and LGB were less than 3 h. However, in each case the onset of the transcriptional reactivation was completed by 8 h *ex vivo* incubation in those cells that expressed detectable HTLV-1 mRNAs (20% to 80% of the total population, again depending on the HTLV-1^+^ subject examined); the remaining cells showed no sign of proviral transcription during the period of observation.

**Fig 3.**
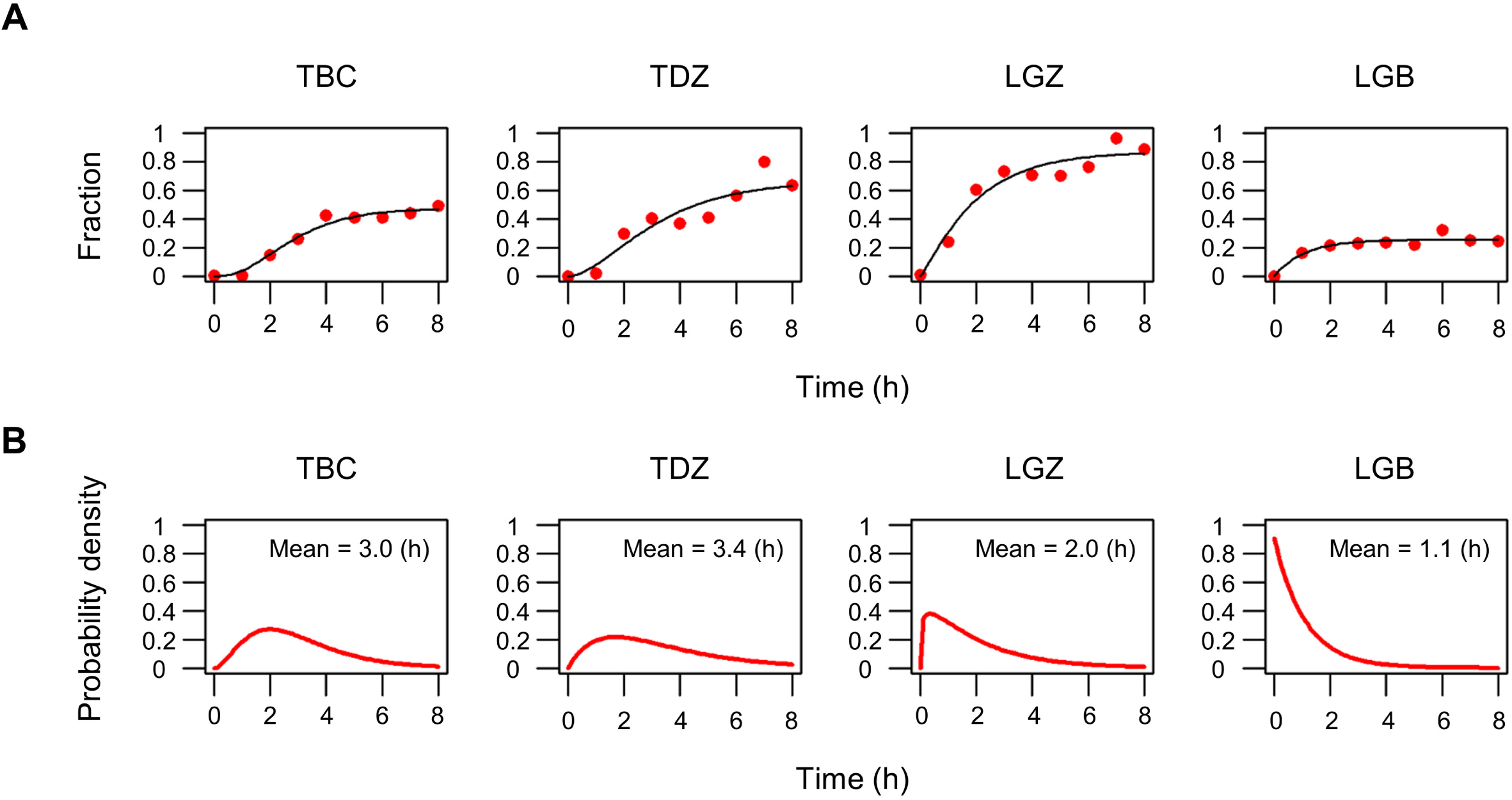
Rate of onset of HTLV-1 reactivation. (A) Increase in the fraction of cells that carry ≥3 sense transcripts over time (red dots). The black line for each HTLV-1^+^ subject indicates a fitted curve with a cumulative gamma distribution. (B) Probability density of the gamma distribution obtained from panel A for each HTLV-1^+^ subject. The inset numbers indicate the mean time of the onset of reactivation after the start of *ex vivo* culture.

### Stochastic simulation predicts a burst termination following the promoter activation

From the results presented above, we established the following characteristics of the transient and heterogeneous HTLV-1 transcription in *ex vivo* PBMCs during the short-term culture. (1) The timing of the onset of transcriptional reactivation peaks in the first few hours of *ex vivo* incubation. (2) The sense-strand transcription is restored with an initially low transcription rate. (3) Subsequently the rate of transcription of the sense strand increases rapidly: we postulate that this rapid sense-strand transcription is triggered by the strong positive feedback loop exerted by Tax protein. (4) Only a subset of HTLV-1^+^ PBMCs (20% to 80%, depending on the subject) restore the sense transcription.

We wished to identify the smallest number of parameters needed to explain the observed kinetics of proviral transcription. We applied a three-state promoter model [21] with some modifications (Fig 4A): the model allows the HTLV-1 sense promoter to transit from the “Off” to “On” state, and further progress to the “On-Tax” state by binding Tax protein; transcription occurs at the On state and the On-Tax state, with distinct transcription rates *k*_5_ and *k*_6_, respectively (*k*_5_ < *k*_6_). We used the smFISH data from the ATL patient LGZ for the parameter optimisation, because the single dominant clone of HTLV-1^+^ PBMCs in this leukemic subject requires only a single kinetic parameter for each reaction rate in the model, as opposed to a polyclonal population in which each clone would require a unique reaction rate to be assigned. The simulations of this three-state model are shown in Fig 4B. The three-state model failed to capture the observed distribution of the transcript count in LGZ (Fig 4B, upper panel): it led to an overestimation of the number of sense-strand transcripts at the later time points. Therefore we assumed an additional reaction path in the three-state model (Fig 4B, lower panel), which allows the promoter to transit directly from the On-Tax state to the Off state with the parameter *k*_10_. This model fitted the smFISH data well (Fig 4B, lower panel; see also S1 Text) and accurately interpolated missing data (S3 Fig); the resulting parameter estimates (posterior distributions) are presented in Fig 4C. The parameter *k*_10_, which is a strong determinant of the duration of the sense-strand transcription, converged within the range of 1 to 3 (h^−1^) (Fig 4C and S4 Fig), indicating that the HTLV-1 sense-strand promoter terminates the Tax-activated rapid transcription in less than an hour (mean duration ~ 1/*k*_10_) in this putatively malignant clone. The duration of the rapid transcription phase was measured in each cell in the simulation by recording the interval between the initiation of the On-Tax promoter state and the direct transition from the On-Tax to the Off state (Fig 4D). These results indicate that the HTLV-1 sense-strand transcription is rapidly terminated.

**Fig 4.**
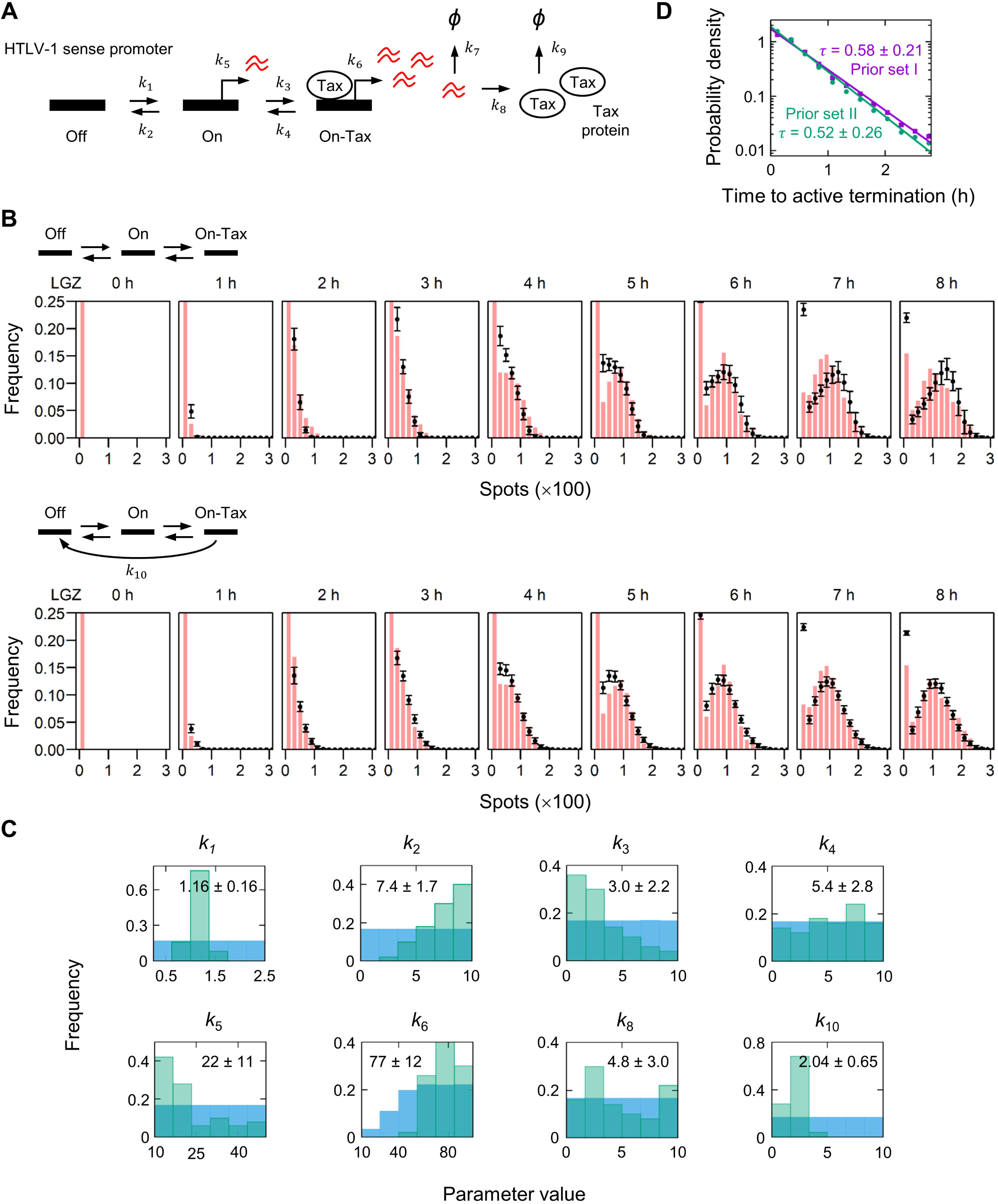
Stochastic simulation of the transient HTLV-1 expression in PBMCs *ex vivo*. (A) Kinetic model of HTLV-1 proviral transcription, with three promoter states: Off, On, and On-Tax. (B) Stochastic simulation of the model presented in panel A. The best 50 fits (mean and standard deviation shown in black) are superimposed on the smFISH data from the ATL patient LGZ (red bars, reproduced from Fig 2B). (C) The marginal distributions of the best 50 parameters (posterior; green bars); the respective prior distributions are shown in blue. The mean and standard deviation are indicated in the inset. (D) The interval between the initiation of the On-Tax promoter state and the direct transition from the On-Tax to the Off state in the simulation. The frequency of the interval was fitted with an exponential decay, and its probability density function is presented. The mean intervals (τ) ± standard deviation for the prior set I (Table 2) and the prior set II (S4 Fig) are indicated in the inset.

### Between 20% and 70% of HTLV-1-infected PBMCs are *HBZ*-negative at a given time

Lastly, we investigated the expression of the HTLV-1 antisense transcript *HBZ*. PBMCs isolated from one asymptomatic HTLV-1^+^ carrier (HBL), three patients with HAM (TBC, TDK and TDZ) and four with ATL (LGZ, LFV, LFK and LGB) were examined by smFISH (Fig 5A). The distribution of the *HBZ* mRNA count per HTLV-1^+^ PBMCs (without incubation) is shown in Fig 5B; the mean (<m>) is indicated in the inset. These results lead to two conclusions. First, the number of *HBZ* mRNA transcripts carried per cell was very low: frequently only one or two molecules of *HBZ* mRNA were present per cell; and unexpectedly 20% to 70% of HTLV-1^+^ PBMCs in the subjects (except LFK) were negative for *HBZ* at a given time. In the subject LFK, almost all the HTLV-1^+^ PBMCs were negative for *HBZ* (also see S2 Fig). Second, the distribution of the number of *HBZ* molecules conformed to the Poisson distribution of the parameter <m>, the observed mean mRNA count (Fig 5B, black lines; the p-value of the Kolmogorov-Smirnov test is indicated in the inset). This observation indicates that the *HBZ* transcripts are produced and degraded at a constant average rate (Fig 5C). Our previous study showed that the half-life of *HBZ* mRNA is 4.4 h [14], from which we estimate the degradation rate δ to be ln2 / 4.4 = 0.1575 h^−1^. Since the mean number of *HBZ* mRNA molecules carried per cell <m> ranged from 0.380 to 1.31 (Fig 5B, except LFK), we estimate that the mean production rate of *HBZ* mRNA (*k* = <m>·δ) falls within the range 0.0599 to 0.206 h^−1^ (Fig 5C).

**Fig 5.**
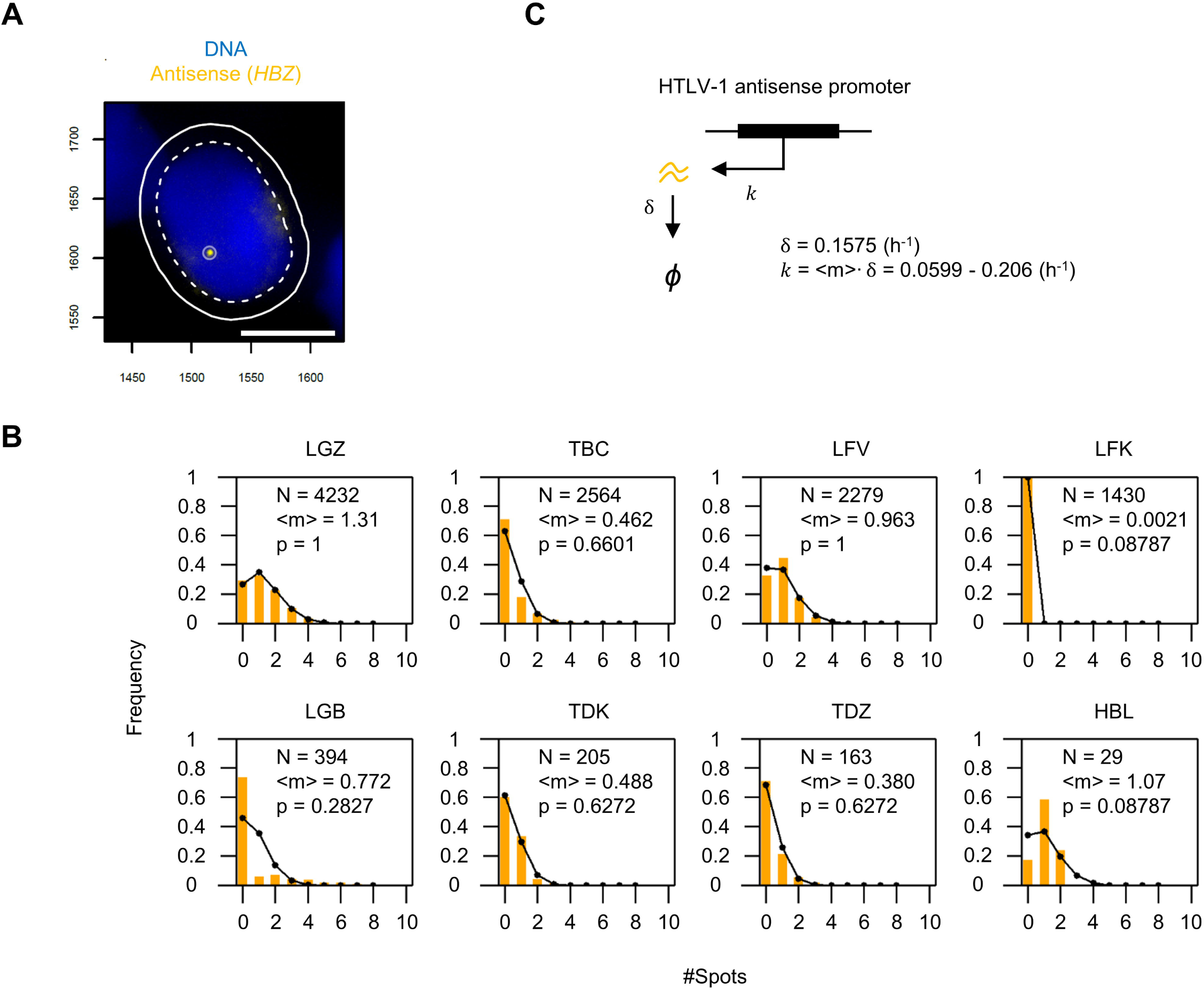
Identification of HTLV-1 antisense transcripts by smFISH. (A) Typical image captured by smFISH of *HBZ* mRNA (yellow spot) in a cell (from a patient with HAM, coded TDZ) without *ex vivo* incubation. Blue indicates the nucleus stained with DAPI. Scale bar (white) = 5 μm. (B) Frequency distribution of the number of *HBZ* mRNA molecules carried per HTLV-1^+^ PBMC. Yellow bars show the frequencies observed by smFISH; the number of HTLV-1^+^ cells analysed (N) and the average number of *HBZ* molecules per HTLV-1^+^ cell (<m>) are indicated in the inset. Black lines indicate the expected frequencies from a Poisson distribution with the parameter <m>. (C) The kinetic model for the antisense transcription. *HBZ* mRNA is transcribed at the rate *k*, and degraded at the rate δ.

## Discussion

It is assumed that the HTLV-1 sense strand is expressed *in vivo* [1], because a strong, chronically activated CTL response against Tax and other sense-strand-encoded proteins is readily detectable in PBMCs from HTLV-1^+^ subjects *ex vivo* [11]. However, in the present study HTLV-1 sense-strand transcripts were rarely detected in fresh PBMCs without incubation, using the sensitive and specific method of smFISH, in any of the HTLV-1^+^ subjects examined (Figs 1B and 2B). The results show that HTLV-1 sense-strand transcription is rare in the circulation. We postulate that HTLV-1 expresses the sense strand in different anatomical compartments of the body other than the blood, such as bone marrow and lymphatic organs. This notion is supported by the observation that *tax* is highly expressed in bone marrow in Japanese macaques infected with simian leukemia virus STLV-1, the simian counterpart of HTLV-1 [22]. The smFISH technique could be applied to analyse specimens from those organs, to identify and localize the subset of cells that express *tax*, and to reveal the structural and spatial distribution of HTLV-1 expression *in situ*.

Whereas the sense-strand transcript *tax* is usually undetectable by reverse-transcription-PCR in PBMCs isolated from HTLV-1^+^ subjects, the HTLV-1 antisense transcript *HBZ* is invariably detected in the same subjects [10] at the cell population level. This observation raised the hypothesis that HBZ protein and *HBZ* mRNA play an essential part in HTLV-1 persistence (reviewed in ref [23]).

In our previous study, we examined *HBZ* expression by smFISH in five HTLV-1-infected T cell clones isolated by limiting dilution from HTLV-1^+^ patients and maintained *in vitro* [14]. While the distribution of the *HBZ* mRNA count in three of the clones showed a small but significant departure from the expectation derived from the Poisson distribution, the other two clones conformed to the Poisson distribution (S5 Fig). The slight deviation from the Poisson distribution observed in certain clones may result from a small burst of *HBZ* transcription [14] (S2 Text). The average *HBZ* mRNA count per cell in those five clones was in a small range (1.67 to 2.92), suggesting that the *HBZ* production rate varies little with the genomic insertion site of the HTLV-1 provirus.

In PBMCs circulating in HTLV-1-infected individuals, we assume that the great majority of HTLV-1^+^ clones are capable of expressing *HBZ*, because its sequence is highly conserved and because *HBZ* appears to be necessary for the persistence of HTLV-1^+^ T lymphocytes *in vivo* [23]. In the *ex vivo* PBMCs, we did not observe strong transcriptional bursts of the antisense strand. The distribution of the *HBZ* count in the patients with ATL conformed well with the Poisson expectation (Fig 5B), except in the sample from the patient coded LGB. This leukemic patient was anomalous in that only 20% of the total population expressed the proviral sense strand, even though a single dominant clone was present in the peripheral blood. The malignant clone present in another ATL patient (LFK), which essentially expressed no *HBZ*, had presumably acquired mutations that allowed the cells to proliferate independently of the HTLV-1 provirus. The *HBZ* count in the patients with HAM, with a polyclonal HTLV-1^+^ T cell population, also conformed to the Poisson distribution. We conclude that *HBZ* is produced constantly in each infected cell, at ~0.1 molecules per hour on average; due to the low average rate of transcription, around half of all HTLV-1-infected cells are *HBZ*-negative at any one time.

It has been reported that *HBZ* mRNA promotes cell proliferation *in vitro* [10, 24]. If this observation holds true in naturally-infected cells, with the low abundance of *HBZ* mRNA molecules observed in this study, it becomes increasingly important to understand how *HBZ* mRNA exerts such functions.

The HBZ protein level is also of great interest because the CD8^+^ T cell response against HBZ is a significant determinant of the HTLV-1 proviral load [25], the main correlate of the risk of HAM and ATL [1]. A previous study from others estimated, by western blotting, that the abundance of endogenous HBZ protein is 17 molecules per cell in an ATL case [26]. In addition to the abundance of *HBZ* mRNA quantified in the present study, measuring the half-life of HBZ protein will permit the estimation of the variance in HBZ protein expression as well as the rate of HBZ translation.

During the *ex vivo* incubation, the distribution of *HBZ* mRNA molecules per cell was stable compared to the changes in the sense-strand expression (S6 Fig) in three of the four subjects examined (TBC, TDZ and LGB). This observation indicates that the antisense strand is not reactivated in concert with the sense strand. In the fourth subject examined (LGZ), the abundance of *HBZ* mRNA in the population decayed over time (S6 Fig, LGZ). Since the majority of HTLV-1^+^ cells in LGZ (80%) restored the sense-strand transcription, it is possible that transcriptional interference - by the collision of two converging polymerase complexes - diminished the antisense transcription in this subject. However, in each of the remaining three subjects studied, there was no correlation between the sense-strand transcription and *HBZ* transcription. These results suggest that transcriptional interference between the sense- and antisense-strands does not play an important part in regulating HTLV-1 proviral transcription in most infected T cell clones.

By applying smFISH to the transient HTLV-1 transcriptional reactivation of the sense strand, we found that reactivation was highly heterogeneous in *ex vivo* PBMCs, as shown by three observations. First, only a subset of the HTLV-1^+^ PBMCs reactivated proviral transcription; second, the time of onset of transcription varied between cells; and third, the intensity of reactivation (i.e. the number of transcripts produced in a given period of time) varied widely in the population. Our results indicate two distinct phases of sense-strand transcription: the initial low-rate transcription produces ~10 to 20 mRNA molecules; thereafter a phase of rapid transcription occurs, probably attributable to the positive feedback exerted by the viral transactivator protein Tax. Transcriptional heterogeneity is likely to arise from the difference between clones in the HTLV-1 proviral integration site in the host genome. However, even in the ATL subjects examined in this study (LGZ and LGB), who carried one dominant clone in which (by definition) each cell shares the same integration site, a significant variance between cells in reactivation intensity was observed, which was equivalent to that of HTLV-1^+^ patients with polyclonal insertions (Fig 2C). Such heterogeneity in transcription is often ascribed to the stochasticity that inevitably arises in biological processes.

In this study, we demonstrated that HTLV-1 sense-strand reactivation in PBMCs occurs in a burst of rapid transcription. But how long does such a burst last in the nucleus? The precise duration of a burst in individual cells cannot be determined by smFISH, which captures only a single timepoint. By contrast, the stochastic simulation of the three-state promoter model (Fig 4B and S1 Text) estimated that the average duration of the intense transcription in PBMCs from a leukemic patient (LGZ) was less than an hour: the model required the HTLV-1 sense-strand promoter to revert from the On-Tax state to the Off. Without invoking this extra parameter, cells did not terminate the transcription during the 8 h of the simulation, so the number of transcripts overshot the observed smFISH data (Fig 4B and S1 Text). These parameters were estimated from a population of PBMCs containing a single dominant clone: other HTLV-1-infected clones may differ in the magnitude of the kinetic parameters, although the fundamental molecular mechanisms are unlikely to vary. Indeed, the putative malignant clone in the patient LGZ might have been selected *in vivo* by the host immune response for rapid termination of sense-strand transcription: *tax* expression is lost in ~50% of cases of ATL, and it is believed that the strong cytotoxic T lymphocyte response to Tax is responsible for this silencing.

Mahgoub et al. reported that the average duration of Tax expression was ~19 hours [27] in an HTLV-1-infected cell line (MT-1) that was established from a patient with ATL, using a GFP reporter system to indicate the production of Tax protein. By contrast, we used uncultured *ex vivo* PBMCs, which carry a single copy of the HTLV-1 provirus in each cell [28], and quantified the HTLV-1 transcripts by smFISH, which allows direct estimation of the gene transcription kinetics. Based on our model, we postulate that the HTLV-1 sense strand, which encodes *tax* and is activated by the strong positive feedback of Tax protein, terminates the transcription after a short period, even if Tax protein is still present in the cell. However, it is possible that repeated sense-strand bursts occur in rapid succession, due to repeated reattachment of transcription complexes to the HTLV-1 promoter, resulting in the longer duration of Tax presence in the cell observed by Mahgoub et al.

Our data, together with inferences from the mathematical model, suggest that active termination of a transcriptional burst is critical in shaping the dynamics of HTLV-1 sense-strand transcription. We examined the transient (non-equilibrium) dynamics of HTLV-1 transcriptional bursting in primary, naturally-infected lymphocytes, whereas others studied Tat transcription from synthetic Tat circuits to analyse the mechanisms that produce bimodality in Tat expression over an extended parameter regime in a steady-state system [29–31].

The mechanism of this rapid termination of the sense-strand burst remains unknown, as in mammalian gene expression in general. The HTLV-1 accessory protein p30 retains *tax* mRNA in the nucleus, which results in a reduction of Tax protein production, leading in turn to reduced viral transcription [32]. However, p30 is unlikely to account for the rapid termination of the HTLV-1 transcriptional burst in PBMCs observed in the present study, because the abundance of Tax protein continues to rise in PBMCs during 8 h *ex vivo* culture [33]. At least three factors can be identified that may contribute to transcriptional silencing. First, epigenetic modifications at the 5 promoter including re-ubiquitination of histone 2A lysine 119 by polycomb repressive complex 1 (PRC1) [5] and methylation (or hydroxymethylation) of cytosine [34]. Second, sequestration of the transcription co-factors CBP/p300 by HBZ protein [35]. Third, a progressive reduction in Tax translation caused by the accumulation of Rex protein [36]. Recent evidence suggests that transcription factors form phase-separated droplets where transcriptional bursts occur, and intrinsically disordered domains of transcription factors are particularly responsible for forming droplets [37, 38]. HTLV-1 Tax protein has a number of binding partners in the cell [39] and forms speckles in the nucleus that colocalize with active transcription spots [40]. These observations raise the possibility that Tax contributes to the formation of phase-separated droplets in the nucleus that serve as the locus for HTLV-1 transcription. External perturbation can disperse phase-separated droplets [38] and might therefore terminate the burst, leading to a refractory period of complete transcriptional silence.

In HIV-1 infection, the potential use of shock-and-kill treatment is still under debate because all approaches used to date have activated only a small fraction of the proviruses in a host. The results presented here demonstrate that the extent of heterogeneity in retrovirus reactivation is diverse in a population of naturally-infected cells, and suggest a quantitative approach for examining the efficacy of the treatment.

## Materials and Methods

### Ethics statement

EDTA-anticoagulated blood from HTLV-1-infected patients was collected at St Mary’s Hospital (Imperial College London, UK). All donors gave written informed consent in accordance with the Declaration of Helsinki to donate blood samples to the Communicable Diseases Research Tissue Bank, approved by the UK National Research Ethics Service (15/SC/0089).

### Patient-derived PBMCs

Information on the HTLV-1^+^ subjects examined in this study (four ATL patients, three HAM patients and one asymptomatic carrier) is available in Table 1. PBMCs were isolated using Histopaque (Sigma, H8889) and washed with PBS. Unless mentioned otherwise, they were stored in liquid nitrogen with FBS/10% DMSO until use.

**Table 1.** HTLV-1^+^ subjects and PBMCs examined in this study.

| Assay # | Patient code | Disease Status | Blood collected | Storage condition | Sampling time (h) | CD8 removal | Proviral load |
| --- | --- | --- | --- | --- | --- | --- | --- |
| 1 | TDZ | HAM | 06 Jan 2015 | Frozen | 0,1,2,3,4,5,6,7,8 | CD8-depleted | 16.4%<sup>a</sup> |
| 2 | TBC | HAM | 18 Sep 2017 | Frozen | 0,1,2,3,4,5,6,7,8 | CD8-depleted | 81.1% |
| 3 | LGZ | ATL | 08 Jul 2014 | Frozen | 0,1,2,3,4,5,6,7,8 | CD8-depleted | 89.7% |
| 4 | HBL | Asymptomatic | 01 Oct 2018 | Fresh<sup>b</sup> | 0 | Total PBMCs | 1.95% |
| 5 | LFK | ATL | 01 Oct 2018 | Fresh<sup>b</sup> | 0 | Total PBMCs | 46.2% |
| 6 | LFV | ATL | 01 Oct 2018 | Fresh<sup>b</sup> | 0 | Total PBMCs | 94.6% |
| 7 | TDK | HAM | 02 Oct 2018 | Fresh<sup>b</sup> | 0 | Total PBMCs | 11.2% |
| 8 | LGB | ATL | 10 Jun 2014 | Frozen | 0,1,2,3,4,5,6,7,8 | CD8-depleted | 88.8% |

<sup>a</sup>measured in the sample collected on 04 Jul 2016
<sup>b</sup>Samples were used immediately after PBMC isolation, without freeze-thawing.

### Short-term culture of patient PBMCs

Patient-derived PBMCs were thawed and washed with PBS (Gibco, 14190) containing 2 mM EDTA and 0.1% FBS. CD8^+^ cells were removed with Dynabeads (Invitrogen, 11147D) to avoid CTL killing against HTLV-1 Tax^+^ cells. Cells were suspended (1 × 10^6^ cells/mL) in RPMI1640 (Sigma, R0883) supplemented with 2 mM L-glutamine (Gibco, 25030-81), penicillin/streptomycin (Gibco, 15070-063) and 10% fetal bovine serum (Gibco, 10500-064). One mL of cell suspension (1 × 10^6^ cells) was aliquoted and dispensed into 5 mL polystyrene round-bottom tubes (Falcon, 352054) to keep the cell density consistent between assays. Cells were incubated in 5% CO_2_ at 37°C until sampled. One aliquot was sampled at each time point (every hour) and subjected to single-molecule RNA FISH.

### Single-molecule RNA FISH

Cells were washed and suspended in PBS at a density of 5-10 × 10^6^ cells/mL. An aliquot (20 μL) of the suspension was spread on to a coverslip that had been coated with poly-L-lysine (Sigma, P8920). Cells were fixed with 2% formaldehyde (Thermo Scientific, 28908) on the coverslip for 15 min. Coverslips were immersed in 70% ethanol and stored at −20°C until hybridization.

Hybridization was performed as previously described [14]. Namely, permeabilized cells were hybridized overnight at 37°C with probes that target either *gag* (HTLV-1 sense strand; labeled with FAM), *tax* (HTLV-1 sense strand; labeled with Quasar 670) or *HBZ* (HTLV-1 antisense strand; labeled with Quasar 570) in hybridization buffer (Stellaris, SMF-HB1-10). Cells were washed on the coverslip twice with wash buffer A (Stellaris, SMF-WA1-60), and then wash buffer B (Stellaris, SMF-WB1-20). DAPI staining was included during the second wash with the buffer A. The cells were embedded in VECTASHIELD mounting medium (Vector Laboratories, H-1000) for imaging.

The coverslips were imaged with an Olympus IX70 inverted widefield microscope with a 100× 1.35NA UPlanApo oil objective lens (Olympus), a Spectra Light Engine illumination source (Lumencor) and an ORCA-Flash 4.0 V2 digital CMOS camera (Hamamatsu).

### Image analysis

Stacked TIFF images containing 54 z-slices with a 300 nm interval were obtained using Micro Manager [41]. Each slice is composed of four channels (DAPI, FAM, Quasar 570 and Quasar 670). The stacked TIFF images were split into the four channels. Cells were segmented using the DAPI channel with an in-house algorithm which identifies the edge of the nucleus by signal intensity (the algorithm is available at DOI 10.17605/OSF.IO/R6Q8T). Dead cells were identified by the abnormal morphology of nucleus and omitted from the analysis. Spots in each of the three channels (FAM, Quasar 570 and Quasar 670) were detected with FISH-QUANT [42]. A threshold for the signal intensity was set to distinguish the mRNA spots from spurious signals: an example is shown in S2 Fig containing the samples that were hybridized and imaged in the same batch - LGZ fixed at 7 h; LFK and LFV fixed without incubation. The image data are available at DOI 10.17605/OSF.IO/R6Q8T (open under CC0 1.0 Universal Licence).

### Proviral load measurement

DNA was extracted from unfixed cells using DNeasy Blood & Tissue Kit (Qiagen, 69504). The reaction of 20 μL volume was set up using ddPCR Supermix (Bio-Rad, 186-3023) containing 900 nM each of forward and reverse primers and 250 nM FAM-labeled probe to detect the HTLV-1 *pX* locus (forward primer, 5′-CTC CTT CCG TTC CAC TCA AC-3′; reverse primer, 5′-GTG GTA GGC CTT GGT TTG AA-3′; probe, 5′-CGC CTA TGA TTT CCG GGC CCT G-3′). The copy number of a host gene *RPP30* was determined using 1 μL of HEX-labeled PrimePCR RPP30 (Bio-Rad, 10031243) in the reaction to calibrate the amount of DNA input. Droplets were generated with QX200 Droplet Generator (Bio-Rad). PCR was performed (95°C for 10 min; 40 cycles of 94°C for 30 sec and 60°C for 1 min; 98°C for 10 min) with C1000 Touch thermal cycler (Bio-Rad). Fluorescent signal was detected using QX200 Droplet Reader (Bio-Rad).

### Half-life estimation of the HTLV-1 sense transcripts

Two HTLV-1 infected T-cell clones [14] were treated with 10 µg/ml Actinomycin D (Sigma, SBR00013). Total RNA was extracted at 0, 3, 6 and 9 hours of treatment using RNeasy Plus Mini Kit (Qiagen, 74134), and was reverse-transcribed to cDNA using Transcriptor First Strand cDNA Synthesis Kit with random primers (Roche, 04897030001). The relative abundance of HTLV-1 sense-strand transcripts at each time point was measured by qPCR with Fast SYBR Green Master Mix (Thermo Fisher Scientific, 4385612) (forward primer, 5′-CCG GCG CTG CTC TCA TCC CGG T-3′; reverse primer, 5′-GGC CGA ACA TAG TCC CCC AGA G-3′ [5]). The half-life of the HTLV-1 sense transcripts was estimated by non-linear regression curve fitting (one phase decay) using GraphPad Prism version 8.2.1 for Windows (GraphPad Software).

### Stochastic simulation of the transient HTLV-1 sense-strand transcription

A Gillespie stochastic simulation algorithm was employed [43]. The molecular species and the schematic diagram of reactions are shown in Fig 4A. We assumed three possible states for the HTLV-1 sense-strand promoter: the “Off” state where essentially no transcription takes place; the “On” state from which sense transcripts are produced at a low transcription rate; and the “On-Tax” state where the rapid transcription takes place due to the strong positive feedback in the presence of Tax protein. The HTLV-1 sense promoter was set at the Off state in each cell with no sense transcripts and Tax protein in the initial condition. In modelling the HTLV-1 transient transcription in the patient LGZ, 20 % of the total cells in the simulation were confined to the Off promoter state (Fig 3A, LGZ). A random parameter search was performed to identify the sets of parameters that fit to the smFISH data (S1 Text). The parameter search space is shown in Table 2. The limits of the parameter search space were set using the data from smFISH experiments on the timing of the onset of transcription (for *k*_1_) and the number of sense transcripts observed prior to the rapid transcription (for *k*_5_ and *k*_3_). The sense-strand mRNA degradation rate (*k*_7_, ln2 / 4.5 = 0.154 h^−1^) was determined experimentally in the present study (S7 Fig). The degradation rate of Tax protein (*k*_9_) was obtained from Rende et al. [44]. One million parameter sets were tested for each respective model. The goodness of fit for each parameter set was assessed by squared deviation summed across the bins in every time point. See also the supporting information (S1 Text); the code for model iii is available at DOI 10.17605/OSF.IO/R6Q8T.

**Table 2.**
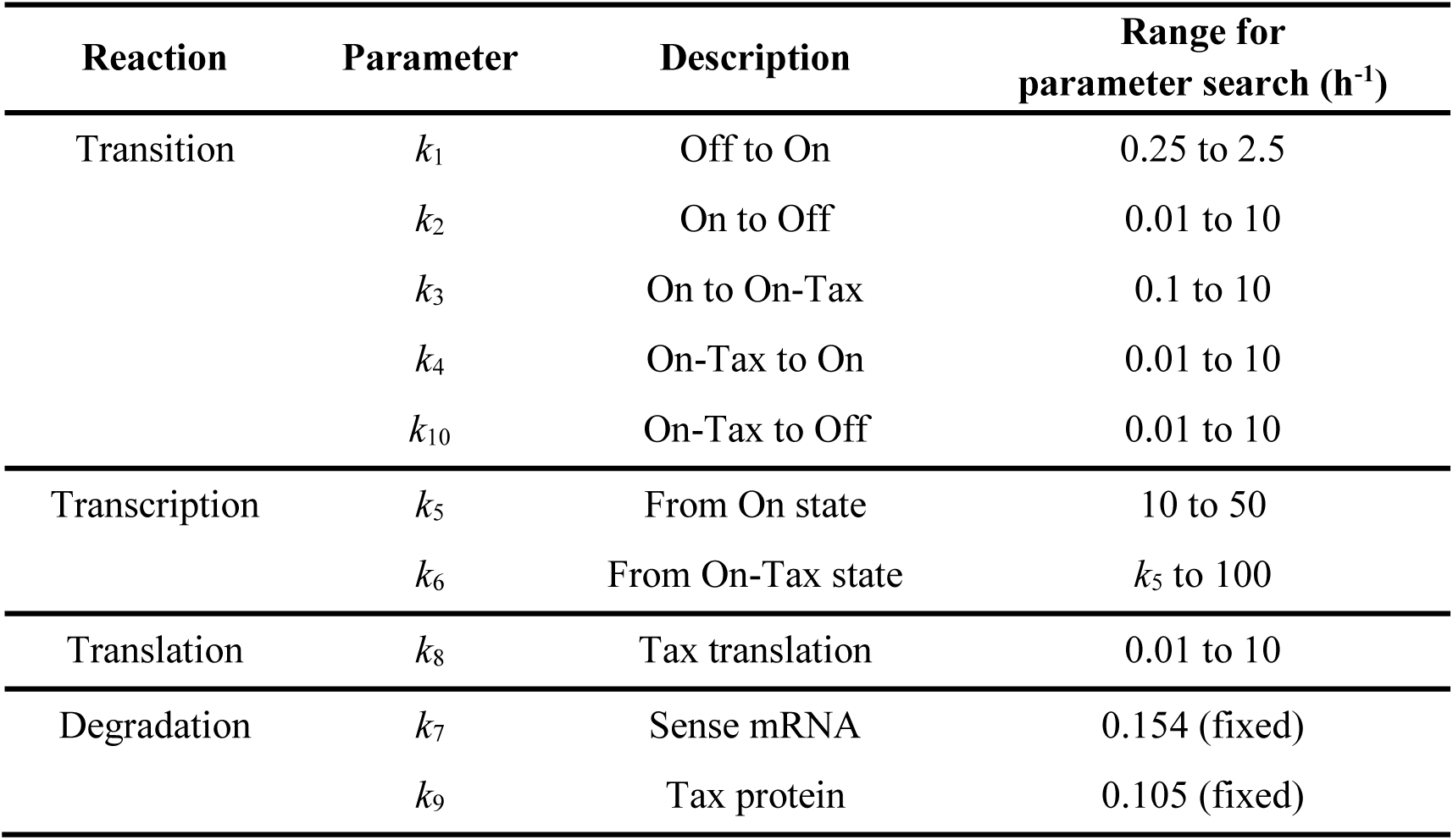
Parameters used in the stochastic simulation.

| Reaction | Parameter | Description | Range for<br>parameter search ($\text{h}^{-1}$) |
| --- | --- | --- | --- |
| Transition | $k_1$ | Off to On | 0.25 to 2.5 |
| | $k_2$ | On to Off | 0.01 to 10 |
| | $k_3$ | On to On-Tax | 0.1 to 10 |
| | $k_4$ | On-Tax to On | 0.01 to 10 |
| | $k_{10}$ | On-Tax to Off | 0.01 to 10 |
| Transcription | $k_5$ | From On state | 10 to 50 |
| | $k_6$ | From On-Tax state | $k_5$ to 100 |
| Translation | $k_8$ | Tax translation | 0.01 to 10 |
| Degradation | $k_7$ | Sense mRNA | 0.154 (fixed) |
| | $k_9$ | Tax protein | 0.105 (fixed) |

## Acknowledgements

We thank Graham Taylor, Lucy Cook, the donors and research nurses in the National Centre for Human Retrovirology, Imperial College; Dr Aileen Rowan for discussion on the ATL subjects; Dr Martin Billman for help with the single-molecule RNA FISH; and the Imperial College Research Computing Service (DOI: 10.14469/hpc/2232) for the use of the high-performance computer cluster; and the Gen Foundation for their support.

## Supporting information

**S1 Fig. Average HTLV-1 sense-strand transcripts in PBMCs in *ex vivo* culture over time.**

**S2 Fig. Determination of the threshold for mRNA spot detection.**

**S3 Fig. Accuracy of data prediction by dynamical models.**

**S4 Fig Parameter estimation.**

**S5 Fig. Antisense *HBZ* expression in *in vitro* maintained HTLV-1^+^ T cell clones.**

**S6 Fig. HTLV-1 antisense expression in PBMCs *ex vivo* culture over time.**

**S7 Fig. Decay-rate of the HTLV-1 sense-strand transcripts.**

**S1 Text. Model Comparison.**

**S2 Text. An alternative model accounting for the departure from the Poisson distribution.**

