## Supplementary material for "Kinetics of HTLV-1 reactivation from latency quantified by single-molecule RNA FISH and stochastic modelling": S1 Fig

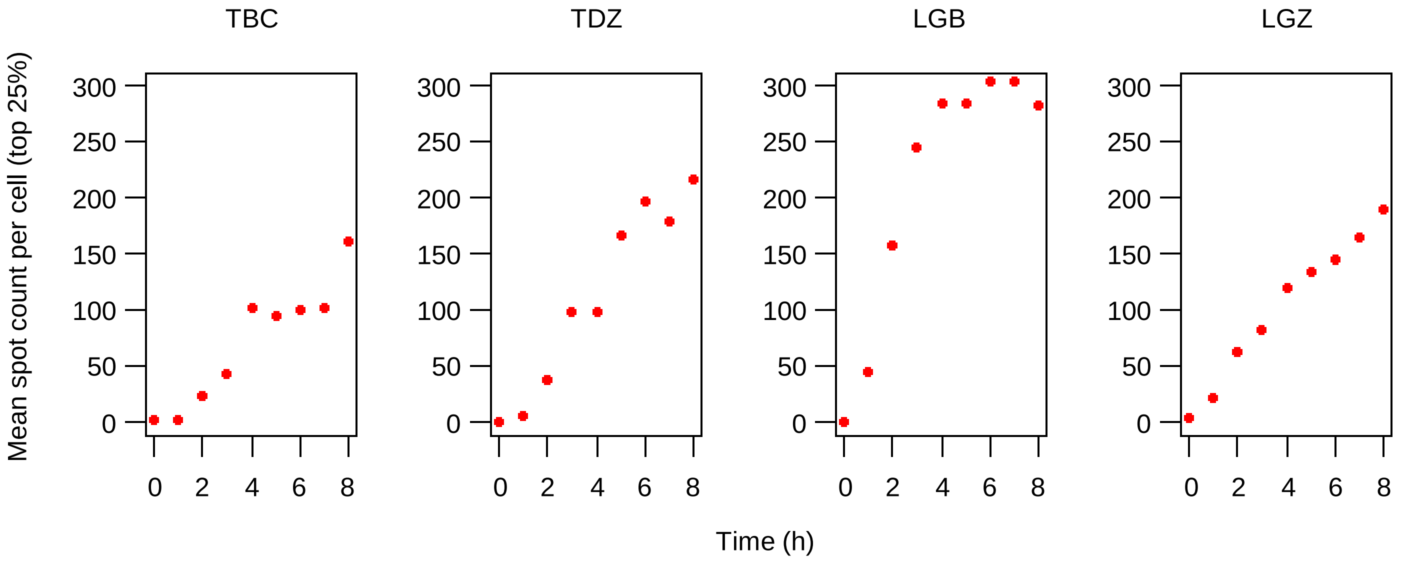


**S1 Fig  Average numbers of HTLV-1 sense-strand transcripts in PBMCs in *ex vivo* culture over time.** The average number of HTLV-1 sense transcripts in the top 25% of all the infected cells at successive timepoints during *in vitro* incubation. The data from two patients with HAM (TBC and TDZ) and two with ATL (LGZ and LGB) are presented.
