## Supplementary material for "Kinetics of HTLV-1 reactivation from latency quantified by single-molecule RNA FISH and stochastic modelling": S2 Fig

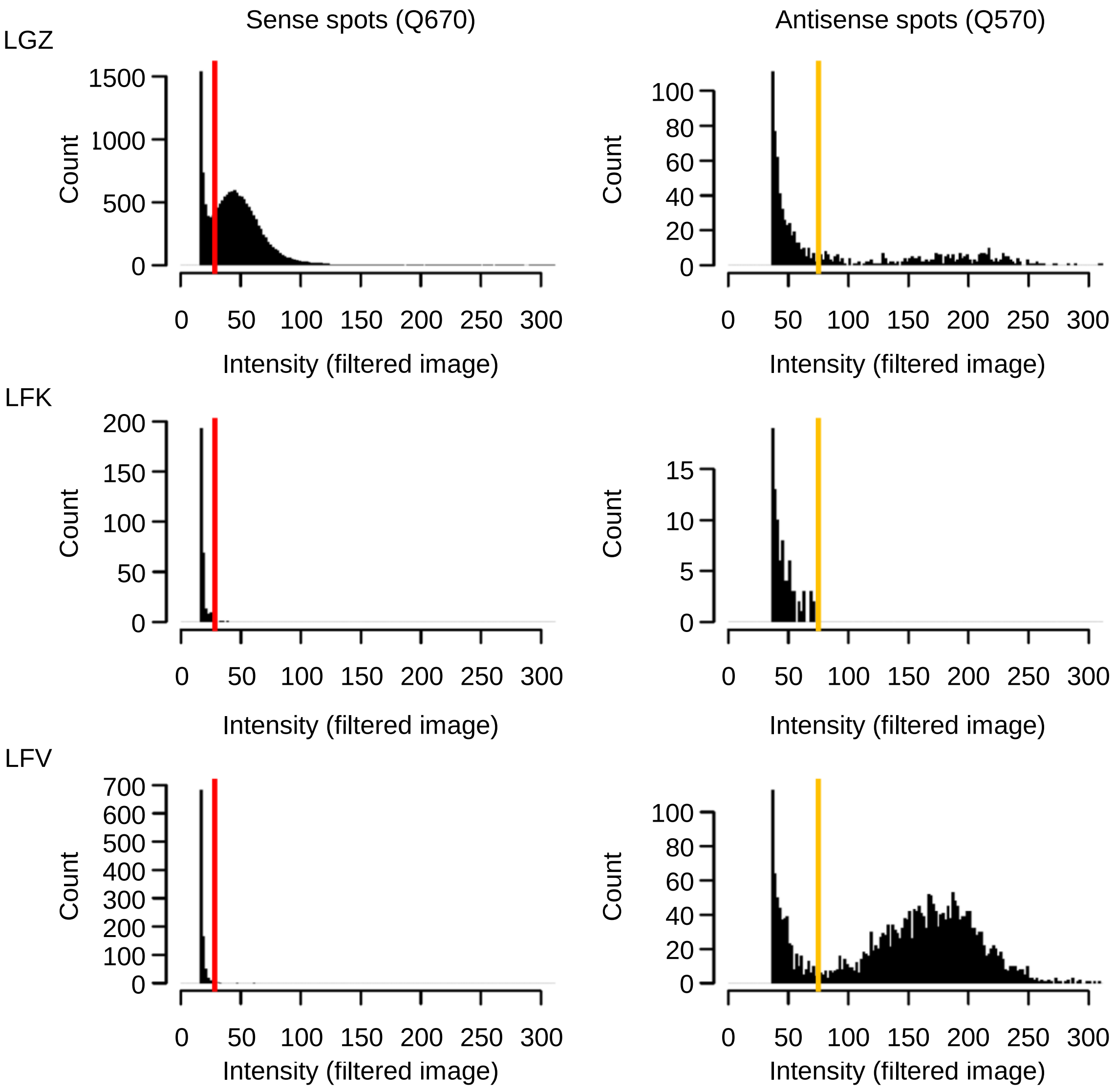


**S2 Fig Determination of the threshold for mRNA spot detection.**

Histograms of the intensity of spots pre-detected with FISH-QUANT (Ref 1) with a conservative threshold setting. The first column shows the spot intensity for Q670, and the second column for Q570. For the analysis, a more stringent threshold was set for each of the channels (red or yellow vertical lines) to remove the spurious signals. The samples shown in the panel (LGZ incubated and fixed at 7 h, LFK and LFV fixed without incubation) were hybridized and imaged in a single batch.
