## Supplementary material for "Kinetics of HTLV-1 reactivation from latency quantified by single-molecule RNA FISH and stochastic modelling": S3 Fig

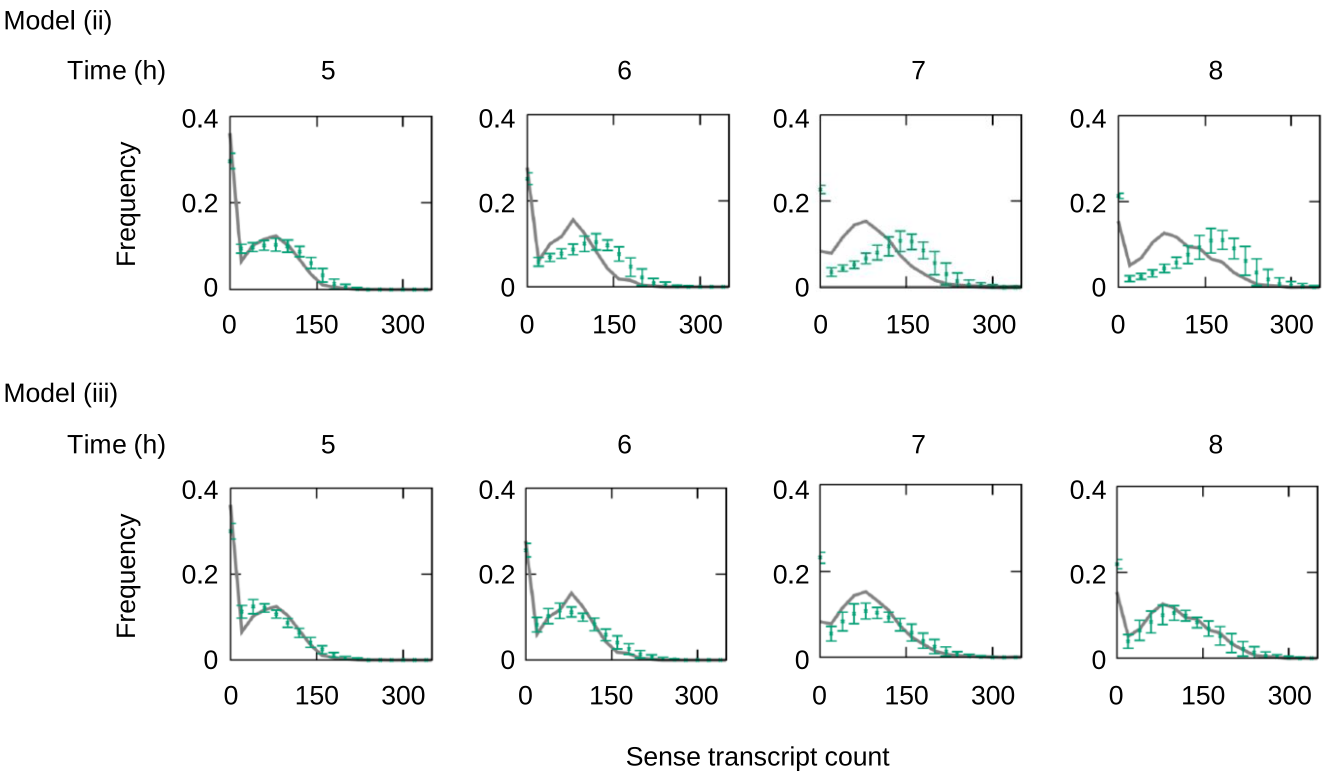


**S3 Fig Accuracy of data prediction by dynamical models.**

To test the performance of the models, we used the experimental data (LGZ) from T = 1, 2, 3 and 4 hrs (omitting the data from T = 5, 6, 7 and 8 hrs) to estimate the best 50 parameter sets from 106 iterations of the models. We used the resulting 50 parameter sets to predict the data points at T = 5, 6, 7 and 8 hrs that were omitted from the parameter estimation. The mean predicted frequency and its standard deviation in each bin are plotted in green, and overlaid on the experimental observation (solid grey line).
