## Supplementary material for "Kinetics of HTLV-1 reactivation from latency quantified by single-molecule RNA FISH and stochastic modelling": S4 Fig

**A**

| Reaction | Parameter | Description | Parameter range ($h^{-1}$) |
|---|---|---|---|
| Transition | $k_1$ | Off to On | 0.1 - 10 |
| | $k_2$ | On to Off | 0.1 - 100 |
| | $k_3$ | On to On-Tax | 0.1 - 10 |
| | $k_4$ | On-Tax to On | 0.1 - 100 |
| | $k_{10}$ | On-Tax to Off | 0.01 - 10 |
| Transcription | $k_5$ | From On state | 0.1 - 25 |
| | $k_6$ | From On-Tax state | 10 - 100 |
| Translation | $k_8$ | Tax translation | 0.01 - 10 |
| Degradation | $k_7$ | Sense mRNA | 0.154 (fixed) |
| | $k_9$ | Tax protein | 0.105 (fixed) (Ref 1) |

**B**

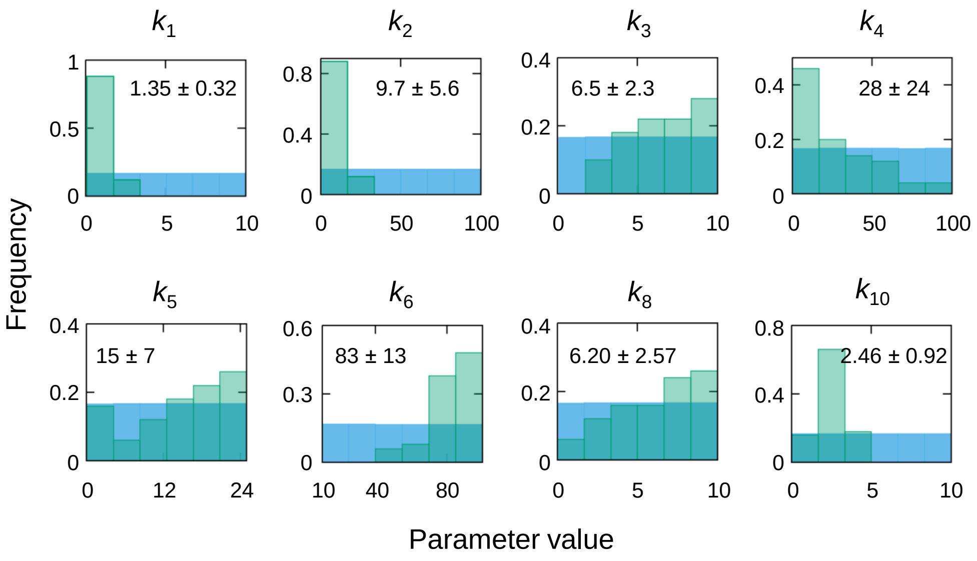


**S4 Fig Parameter estimation.**

(A) To check the parameter convergence, we repeated the parameter search within a second set of prior distributions. (B) We performed $10^6$ iterations with the above prior set and selected the best 50 parameter sets with the minimum distance function (see also S1 Text). The histograms depict the distributions of 50 parameters (green bars) for each reaction in the model; the respective prior distributions are shown in blue. The mean and the standard deviation are indicated in the inset.
