## Supplementary material for "Kinetics of HTLV-1 reactivation from latency quantified by single-molecule RNA FISH and stochastic modelling": S5 Fig

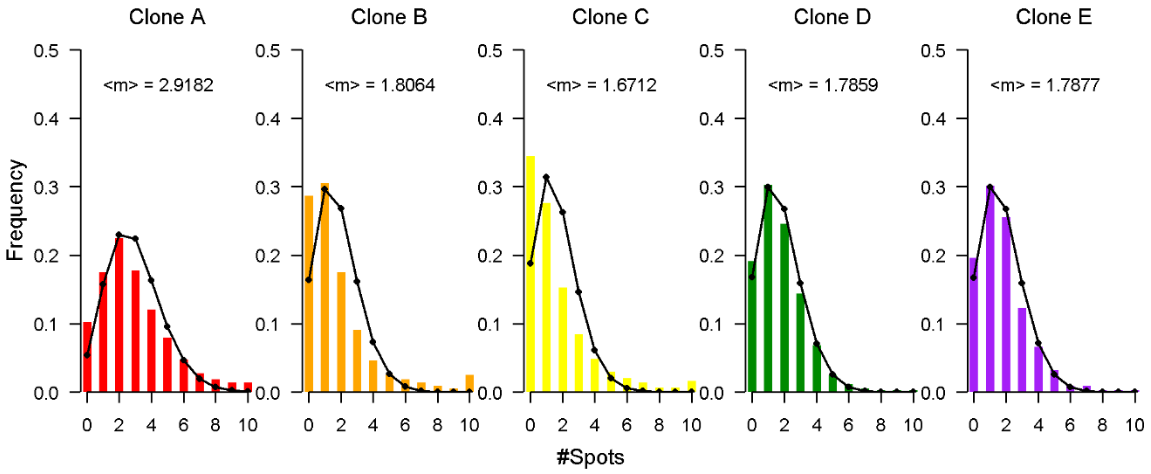


**S5 Fig Antisense *HBZ* expression in *in vitro* maintained HTLV-1⁺ T cell clones.**

The distribution of the number of *HBZ* molecules per cell in *in vitro* maintained HTLV-1$^{+}$ T cell clones. Histograms are reproduced from the graphs presented in Billman *et al* 2017 (Ref 1). The mean number of *HBZ* molecules (<m>) is indicated in the inset. (The last bin is indicated as 10 or more spots in the original paper (Ref 1); in the present figure the bin is shown as =10 spots.) The black line indicates the Poisson distribution with the parameter <m>, the observed mean number of *HBZ* molecules.
