## Supplementary material for "Kinetics of HTLV-1 reactivation from latency quantified by single-molecule RNA FISH and stochastic modelling": S6 Fig

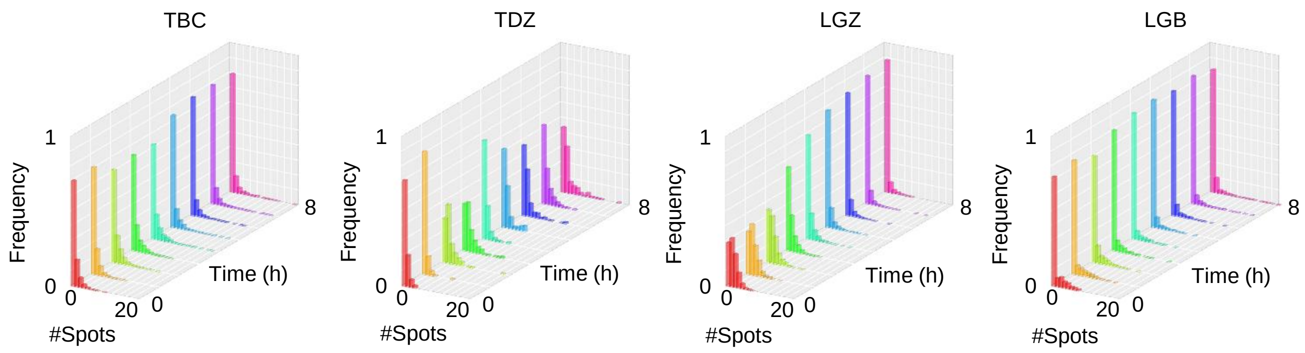


**S6 Fig HTLV-1 antisense expression in PBMCs in *ex vivo* culture over time.**

*HBZ* mRNA count per HTLV-1$^{+}$ cell at successive timepoints during *in vitro* incubation. The data from two patients with HAM (TBC and TDZ) and two with ATL (LGZ and LGB) are presented.
