## Supplementary material for "Kinetics of HTLV-1 reactivation from latency quantified by single-molecule RNA FISH and stochastic modelling": S7 Fig

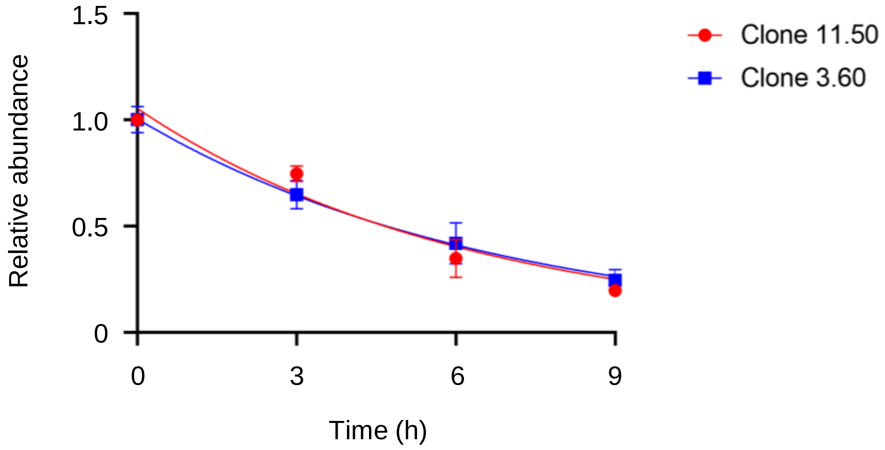


**S7 Fig   Decay-rate of the HTLV-1 sense-strand transcripts.**

Two *in vitro* HTLV-1-infected T cell clones were treated with Actinomycin D to block the transcription, and the abundance of HTLV-1 sense-strand transcripts was quantified at each time point (0, 3, 6 and 9 hours) (see Materials and Methods). The error bars indicate the standard deviation of the qPCR technical replicates. The half-life of the HTLV-1 sense-strand transcripts was estimated to be 4.33 h (Clone 11.50) and 4.66 h (Clone 3.60). The combined estimation of the half-life was 4.45 h, with the 95% confidence interval of 2.70 - 5.61 h.
