## Supplementary material for "Kinetics of HTLV-1 reactivation from latency quantified by single-molecule RNA FISH and stochastic modelling": S1 Text

#### Model comparison

For the parameter estimation, we used the Approximate Bayesian Computation (ABC) method (Ref 1). The experimental data used to fit the model are the frequency distribution of sense transcripts observed at time $T$, which we denote $P_{obs}(m,T)$. The transcript count distributions were binned into intervals of equal size (bin size = 20). For each simulation, we computed the sense transcript distribution at time $T$, $P_{model,\theta}(m,T)$, for a given parameter set $\theta$ randomly drawn from a given uniform prior distribution, as set out in Table 2 in the main text. For the parameter set $\theta$, the following distance function quantifies the mismatch between the experimental observations and the results of simulation:

$$d_{obs,model}(\theta) = \sum_{T}\sum_{i}[P_{obs}(m_i,T) - P_{model,\theta}(m_i,T)]^2$$

where $m_i$ corresponds to the sense transcript count in the *i*-th bin. For each candidate parameter set $\theta^*$, if the distance function $d_{obs,model}(\theta^*)$ is less than a predefined threshold $\epsilon$, we accept the parameter set; otherwise the parameter set is rejected. This is known as the ABC rejection method.

For model comparison, we used the technique proposed in the ABC framework (Ref 1). We inferred the best parameters for the three potential models: (i) two-state model with positive feedback, (ii) three-state model with positive feedback, and (iii) three-state model with active termination of On-Tax (Fig 4B in the main text). Note that $k_1$, $k_2$ and $k_{10}$ were set to 0 for model (i), and $k_{10}$ was set to 0 for model (ii). The simulation was carried out with our codes (written in C++) for 2000 cells, of which 80% were transcriptionally active (Fig 3A, LGZ). For each model, we chose $10^6$ parameter sets from their corresponding uniform priors and calculated the distance function. Let us denote the numbers of parameter sets that satisfy $d_{obs,model}(\theta^*) < \epsilon$ as $N_{\epsilon,model}$. In the ABC framework, the penalty for increasing the number of parameters lies in the fact that the probability of finding a parameter set $\theta^*$ that satisfies the above condition becomes smaller as the number of parameters increases. Assuming the prior probability of choosing a model is uniform, the Bayes factor is the ratio of the posterior probabilities of the two models and provides a measure of the relative goodness of fit of two competing models. In the ABC framework, the Bayes factor for model (i) against model (ii) can be approximated as $B_{i,ii} \approx \frac{N_{\epsilon,i}}{N_{\epsilon,ii}}$ (Ref 1). The table below summarizes the comparisons of the three models. Based on the Bayes factors, we conclude that model (iii) is strongly preferred to the other two.

| Threshold ($\epsilon$) | $N_\epsilon$, after $10^6$ random parameter search from a given prior | | | Bayes factor | | |
|---|---|---|---|---|---|---|
| | Model (i) | Model (ii) | Model (iii) | $B_{iii,ii} \approx \frac{N_{\epsilon,iii}}{N_{\epsilon,ii}}$ | $B_{iii,i} \approx \frac{N_{\epsilon,iii}}{N_{\epsilon,i}}$ | $B_{ii,i} \approx \frac{N_{\epsilon,ii}}{N_{\epsilon,i}}$ |
| 0.06 | 0 | 0 | 1420 | ∞ | ∞ | ∞ |
| 0.1 | 0 | 293 | 16257 | 55.5 | ∞ | ∞ |
| 0.2 | 0 | 17406 | 78926 | 4.5344 | ∞ | ∞ |

### S1 Text

Confirming the result of the model comparison using the ABC scheme, the distance function $d_{obs,model}(\theta^*)$ for the best 50 parameter sets from model (iii) was approximately one-half of that from model (ii), and one-tenth of that from model (i):

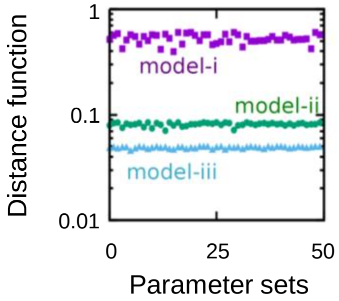


With those 50 parameters for each model, the number of sense-strand transcripts produced in the cells over time was computed. The green dots and bars indicate the mean frequency distribution and its standard deviation from the 50 parameter sets; the observed smFISH data are shown by the solid grey line.

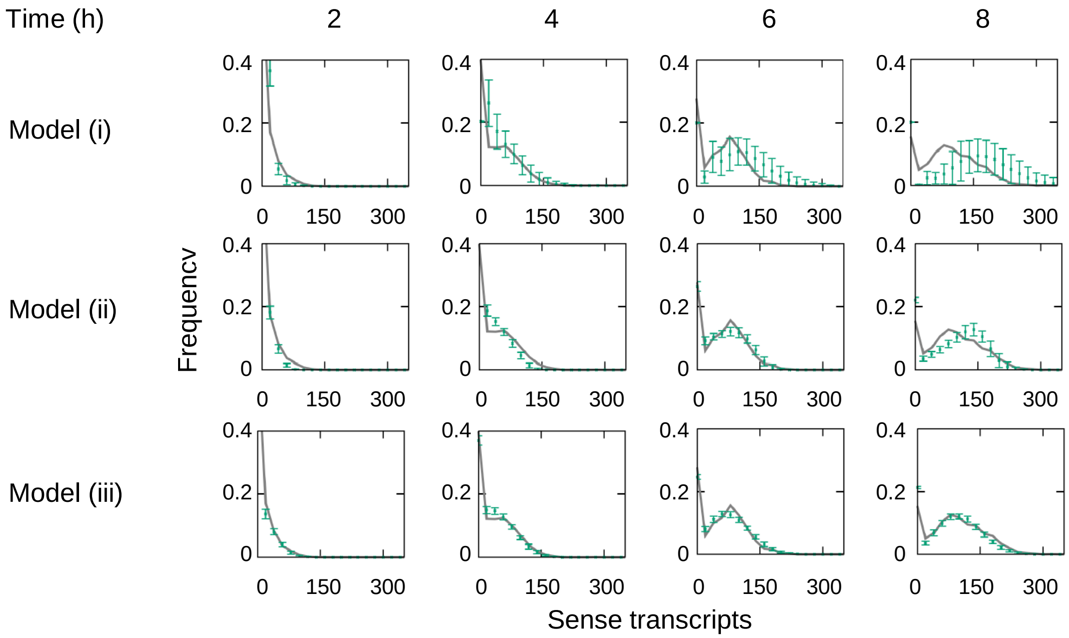


The models (i) and (ii) failed to fit the experimental data (top and middle), whereas the model (iii) reproduced the experimental data accurately. We conclude that both the positive-feedback for the sense-strand transcription and its active termination are required.
