## Supplementary material for "Kinetics of HTLV-1 reactivation from latency quantified by single-molecule RNA FISH and stochastic modelling": S2 Text

### An alternative model accounting for the departure from the Poisson distribution

We observed a deviation in the *HBZ* count per cell in some of the *in vitro* maintained HTLV-1$^+$ T cell clones from the value calculated from the Poisson distribution (Ref 1). Specifically, there was a greater number of *HBZ*-negative cells in each clone at a given time than predicted by the Poisson distribution. We suspected that this deviation was due to the transcription of the antisense strand in small and occasional bursts. Therefore, we applied the following two-state model to clone B as an example.

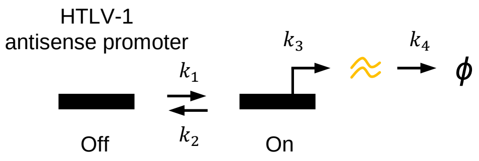


The number of *HBZ* molecules <m> at the steady state can be described as

$$< m > = \frac{k_3[On]}{k_4} = \frac{k_3}{k_4} \cdot \frac{k_1}{k_1 + k_2}$$

We estimated the degradation rate $k_4$ to be 0.1575 h$^{-1}$ as previously reported from experimental measurements (Ref 1). We carried out the parameter search for $k_1$ and $k_3$ within the range of 0.001 to 10 and 0.05 to 50, respectively. The parameter $k_2$ was constrained by

$$k_2 = \frac{k_1 k_3}{< m > \cdot k_4} - k_1$$

We obtained a parameter set ($k_1$=0.658, $k_2$=94.524, $k_3$=41.14 and $k_4$=0.158) which yielded the distribution indicated below by the black line.

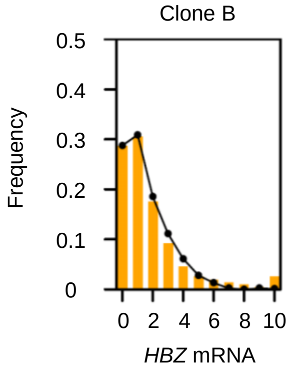


With this parameter set, we estimate the burst frequency and intensity as follows:

burst frequency = $k_1$[Off] = $k_1 \cdot k_2$ / ($k_1$+$k_2$) ~ 0.653 (h$^{-1}$)

burst intensity = $k_3$ / $k_2$ ~ 0.435 (molecules / burst)
